# Antibody-Initiated Loop-Mediated Isothermal Nucleic Acid Amplification (ai-LAMP) as a New Biosensor for Antigen Detection

**DOI:** 10.1101/2025.09.24.678293

**Authors:** Navaporn Sritong, Ana L. Claure Dips, Eric Tan, Karin F. K. Ejendal, Tamara L. Kinzer-Ursem, Jacqueline C. Linnes

## Abstract

Highly sensitive protein detection is critical in research and healthcare diagnostics but remains limited to resource intensive environments. In contrast, lateral flow immunoassay (LFIA)-based diagnostics are popular point-of-care tests due to their simplicity and user-friendly format but lack sensitivity for detecting the low concentrations of proteins that are present early in infections and among asymptomatic individuals. To overcome these limitations, we developed a novel biosensor platform utilizing antibody-initiated loop-mediated isothermal amplification (ai-LAMP) with an LFIA readout for protein detection. This platform integrates protein-specific antibody-antigen binding with the robust signal amplification of LAMP. By conjugating pairs of antibodies to overlapping DNA strands, the presence of a target protein brings the DNA strands into proximity, completing a DNA target to directly initiate the LAMP reaction without any additional ligation step. This approach facilitates the rapid detection of low concentration proteins with a clear visual readout and can be performed in 3 steps from sample to answer. Using ai-LAMP we detected HIV-1 p24 on commercially available LFIAs at a limit of detection (LOD) of 20 fM (0.53 pg/mL), demonstrating 46x improvement over existing HIV p24 LFIAs. The use of ai-LAMP eliminates the need for sophisticated laboratory equipment to detect protein targets in low concentrations, paving a new way for rapid and accessible biomarker detection in clinical and research settings.

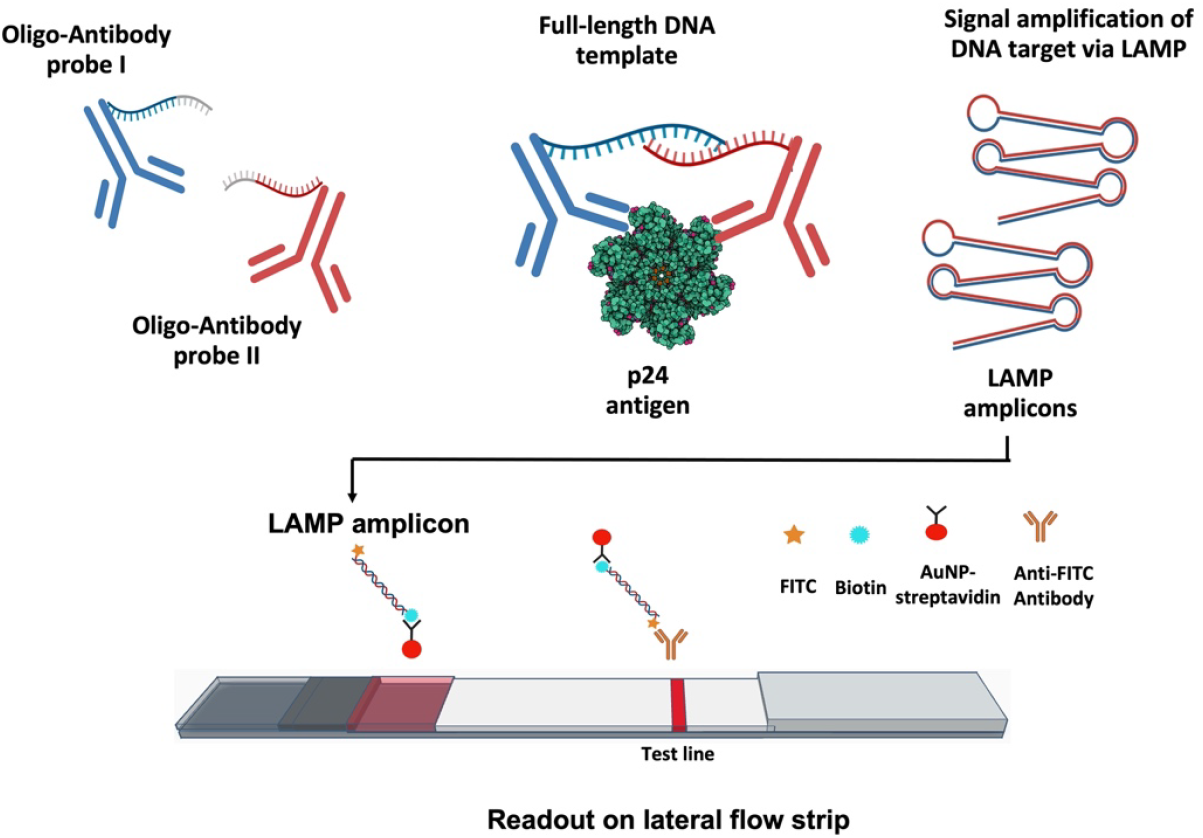

## I. Introduction

Biosensors have emerged as essential tools in medical diagnostics, offering rapid, sensitive, and specific detection of biomarkers. These are especially important in point-of-care (POC) settings, where resources are limited, and quick accurate results are critical for effective disease diagnosis and timely results are essential for clinical decision-making. Lateral flow immunoassays (LFIAs) have become indispensable tools at the POC, offering user-friendly, portable, and cost-effective platforms for on-site diagnostics. However, current LFIAs lack the sensitivity required to detect low-abundance proteins, which can lead to false negatives and missed early diagnoses of infections. While nucleic acid amplification can produce billions of copies of low-abundance targets, translating similar amplification strategies to proteins remains a significant challenge due to the absence of rapid protein amplification mechanisms.

Unlike nucleic acid targets, which can be amplified by polymerase chain reaction (PCR) or other enzymatic methods to achieve ultra-sensitive detection, proteins cannot be directly amplified in a similar manner. To address this challenge, proximity-initiated nucleic acid amplification techniques such as proximity ligation assay (PLA) and proximity extension assay (PEA) have been developed and can yield protein detection at femtomolar (fM) concentrations [1][2]. These methods use pairs of DNA-conjugated antibodies that bind to nearby epitopes on the target protein, enabling the amplification of completed DNA sequence created by the pairs. Typically this amplification is performed using thermal cycled PCR with a fluorescence readout [3]. An alternative isothermal amplification method, rolling circle amplification (RCA), has been explored to eliminate the need for thermal cycling, but RCA suffers from slow amplification and the assay still requires fluorescent detection instrumentation [4].

To overcome the limitations of existing PLA and PEA methods, we develop a novel approach that integrates loop-mediated isothermal amplification (LAMP) with an LFIA readout for rapid, ultrasensitive detection of proteins with minimal equipment. Using the HIV p24 protein as a model target, we first designed orthogonal LAMP primers and DNA templates to avoid cross-reactivity with both HIV and human nucleic acid sequences. We then used Click chemistry to conjugate two partially overlapping DNA strands to pairs of anti-p24 antibodies for oligo-antibody probes, which facilitates high-specificity binding and recognition of the p24 antigen. The presence of a target protein facilitates the hybridization of a complete DNA template, initiating the LAMP reaction. The results are visualized on an LFIA by tagging the LAMP primers, permitting amplicons to bind the LFIA and allowing for clear differentiation between positive and negative results without the need for sophisticated laboratory equipment.

Our assay achieves a limit of detection (LOD) of 20 fM (0.53 pg/mL) for the p24 antigen, demonstrating significant enhancement in sensitivity compared to conventional LFIAs. By combining the strengths of LAMP with an LFIA readout, this platform offers a powerful solution that could be adapted for ultrasensitive at the point of care.

## 2. Material and methods

### 2.1 Reagents

Monoclonal antibodies (mAbs) specific to HIV-1 p24 protein included mAB247p, and mAB249p were purchased from BBI solution (Cardiff, United Kingdom). HIV-1 p24 antigen was purchased from Sino Biological US Inc (Wayne, PA). Primers, ssDNA target with 5’ amine modification, and 1X IDTE (1X TE solution) buffer were ordered from Integrated DNA Technologies (Coralville, IA). Reagents for LAMP reactions included Bst 2.0 DNA polymerase and isothermal amplification buffer from NEB (Ipswich, MA), EvaGreen from VWR International (Radnor, PA), ROX reference dye from Thermo Fisher Scientific (Waltham, MA), and nuclease-free water from Invitrogen (Waltham, MA). The click chemistry reagents including dibenzocyclooctyne-PEG4-N-hydroxysuccinimidyl ester (DBCO-PEG4-NHS), Azido-PEG4-NHS Ester were purchased from Vector Laboratories (Newark, CA). Zeba™ spin desalting columns with 7K molecular weight cut-off (MWCO) and a volume of 0.5 mL were obtained from Thermo Fisher Scientific (Waltham, MA). Amicon® ultra centrifugal filter column with 10 kDa, 50 kDa, and 100 kDa MWCO with a volume of 0.5 mL were obtained from Millipore Sigma (Darmstadt, Germany). The reagents for SDS-PAGE gel electrophoresis included 4–20% Mini-PROTEAN® TGX™ Precast Protein Gels and 4x Laemmli sample buffer, and Precision Plus Protein™ Kaleidoscope™ from Bio-rad (Hercules, CA), Fast DNA Ladder from NEB (Ipswich, MA), SYBR Gold staining from Thermo fisher scientific (Wilmington, DE), and Instant Blue Coomassie protein stain from Abcam (Cambridge, United Kingdom). SARS-CoV-2 Spike Protein was purchased from Sino Biological US Inc., (Wayne, PA)

### 2.2 LAMP primer and template design

A primer set targeting cholera enterotoxin subunit A *(ctxA)* gene of *Vibrio cholerae* from a previously published article [5] was selected as our primer for LAMP reaction as seen in Table S1. The corresponding target sequence of 193 nucleotides (nt) was chosen as the full-length DNA target, which serves as a positive control of LAMP reaction. The full-length target sequence was separated into partial forward and reverse targets; 112F and 125R with small overlapping complementary regions of 6 nucleotides as shown in Table S2. Three LAMP assays were performed to ensure that two partial DNA targets—112F and 125R— are essential for LAMP reaction without false-positive amplification. The LAMP assay components were displayed in Table S3. The experiments included the LAMP reaction with the presence of either 112F or 125R in various concentrations and the mixture of 112F and 125R as templates. In these experiments, the synthesized full-length of 193 bp was used as a positive control while nuclease-free water was used as no template control (NTC). Each type of DNA template was diluted with PBS buffer in 10-fold serial dilution manner from a stock solution of 100 μM to achieve concentrations ranging from 1 pM to 100 nM. The reactions were run at 66 °C for 60 minutes by using the QuantStudioTM 5 Real-Time PCR Instrument (Applied Biosystems, Waltham, MA). The amplification process was monitored by real-time fluorescence data of EvaGreen intercalating dye and ROX reference dye over duration of 60 minutes.

### 2.3 Antibody-oligonucleotide conjugation

The pair of mAbs used in this work were mAB247p (47p) and mAB249p (49p). The two partial DNA targets, 112F and 125R, were conjugated to 47p and 49p, respectively via copper-free click chemistry. The antibodies were activated with azido-PEG4-NHS ester, while the 5’ amine modified oligonucleotides were activated with DBCO-PEG4-NHS. We followed the published conjugation protocols from Iivari Kleino et al. [6] and Wiener et al. [7] with some modifications as noted below. The conjugation process consisted of four major steps as follow:

#### Antibody activation with azido-PEG4-NHS

The azido-PEG4-NHS ester was dissolved in DMSO and diluted to attain a concentration of 10 mM. Then, a 100 μL of 1.2 mg/mL or 8 μM antibody was transferred to a low-protein-binding microcentrifuge tube, followed by the addition of 10 mM DBCO-PEG4-NHS to achieve five molar excess of azido-PEG4-NHS to antibody and incubation at room temperature for 60 minutes under rotary shaking. To quench the NHS present in the free azido-PEG4-NHS, 1/10th of the reaction volume of 1 M tris pH 8.0 was added and incubated at room temperature for 5 minutes. After the antibody-cross-linker reaction, a 7 kDa molecular weight cutoff Zeba desalting column (Thermo Fisher Scientific, Wilmington, DE) was employed to remove the linker. To verify the concentration of azide-activated antibody, the absorbance at 280 nm was recorded by a Nanodrop spectrophotometer (Thermo Fisher Scientific, Wilmington, DE).

#### Oligonucleotide activation with DBCO-PEG4-NHS

DBCO-PEG4-NHS was dissolved in anhydrous dimethyl sulfoxide (DMSO) and was diluted with DMSO to achieve final concentration of 10 mM. The 5’ amine modified 112F and 125R were dissolved in the manufacturer’s 1x IDTE at a concentration of 100 µM. The DBCO-PEG4-NHS in DMSO was added to each aqueous oligonucleotide solution to achieve a 5 molar excess of DBCO-PEG4-NHS relative to the oligonucleotides. The activation was incubated at room temperature for 2 hours under agitation. To quench the NHS present in the free azido-PEG4-NHS\, 1/10th of the reaction volume of 1 M tris pH 8.0 was added and incubated at room temperature for 5 minutes. Excess DBCO-PEG4-NHS was eliminated through ultrafiltration spin filters with a 10 kDa MWCO. The concentration of the DBCO-modified oligonucleotide was determined through absorbance spectroscopy at 260 nm by Nanodrop spectrophotometer.

#### Conjugation of azide-activated antibody and DBCO-activated oligonucleotides

The DBCO-modified oligonucleotides were combined with azide-activated antibodies to achieve a five-fold molar excess of oligonucleotide to antibody (in a 5:1 molar ratio oligonucleotide to antibody). The mixture was incubated overnight (16 – 18 hours) at 4 °C. The conjugate products were purified utilizing Amicon® ultrafiltration columns with 100 kDa MWCO spin filters. To mitigate losses arising from antibody adherence to the filter, the filter columns were treated with a 1% BSA solution in PBS buffer after standard PBS washing. After loading the conjugated antibody into the spin column, the washing process was repeated three times by using 1x PBS pH 7.4. Throughout the washing steps, the speed of centrifugation was set at 12,000 x g for 3 minutes. The purified oligo-conjugated antibody was collected by spinning the filter/spin column at 1000 × g for 2 minutes upside down and transferred to a clean low-protein-binding tube. The concentration of eluted oligo-conjugated antibody was measured by Nanodrop spectrophotometer at a 280 nm absorbance.

#### Analysis of oligo-conjugated antibody probes by SDS-PAGE

The conjugate products were confirmed through non-reducing SDS-PAGE gel electrophoresis by using 4x Laemmli sample buffer. One part of sample buffer was combined with three parts of the sample, followed by heating the mixtures at 95°C for 5 minutes. Precision Plus Protein™ Kaleidoscope™ and Fast DNA Ladder served as the protein and DNA ladders, respectively. The heated samples were loaded to 4–20% precast polyacrylamide gel. The samples were subjected to electrophoresis at 170 V for a duration of 50 minutes in 1X TGS buffer. SYBR Gold was utilized for DNA staining, whereas Instant Blue was employed for protein staining. Post-electrophoresis, the gel was rinsed with deionized water and subsequently stained in 50 mL of 1X TAE containing 5 μL of SYBR Gold, ensuring complete coverage for a duration of 1-1.5 hours to facilitate DNA staining. Gel imaging was conducted under UV302, with the exposure time adjusted to achieve optimal resolution within the range of 40-55 seconds. Following this, the gel was subjected to protein staining in Instant Blue for a duration of 1 hour followed by destaining in deionized water for 2 hours or overnight. Finally, gel imaging under visible light was performed. Band intensities corresponding to unconjugated antibodies and conjugated products on protein-stained gels were quantified using the gels analysis tool in Fiji (ImageJ). Images were preprocessed by automatic adjustment of brightness and contrast, followed by background subtraction with a 50-pixel rolling ball radius. The conjugation yield was determined by calculating the ratio of the conjugate band intensity to the sum of all band intensities.

### 2.4 ai-LAMP assay optimization

#### The optimization of oligo-antibody probes concentrations

To determine the working concentration of conjugate probes, LAMP assays employing a selected primer set were conducted, utilizing oligo-conjugated antibody probes as LAMP targets. Three distinct LAMP assays were performed. The first two assays used either forward oligo-antibody probe (47p112F) or reverse oligo-antibody probe (49p125R) individually across concentrations ranging from 1 pM to 100 nM, which were serially diluted in a tenfold manner with 1X PBS pH 7.4 as templates to mitigate the risk of false-positive amplification attributed to a single type of oligo-conjugated antibody probe. The third assay employed both 47p112F and 49p125R in the molar ratio 1:1 across concentrations ranging from 1 pM to 100 nM as template. Each 25 μL reaction mixture comprised 20 μL of LAMP master mix, which consisted of a 10x primer mix, isothermal amplification buffer, 20X EvaGreen, 50X ROX reference dye, and Bst 2.0 DNA polymerase enzyme and 5 μL of template. The comprehensive details of the full LAMP protocol are outlined in Table S3. For assays utilizing either 47p112F or 49p125R alone as templates, the sample consisted of 2 μL of the respective probe and 3 μL of 1X PBS, while for assays employing both probes, the sample comprised 2 μL of each probe and 1 μL of 1X PBS. Positive controls DNA was full-length DNA target of 193 nt at a concentration of 10 nM, whereas nuclease-free water served as the no template control (NTC). Reactions were conducted at 66 °C for 60 minutes utilizing the QuantStudioTM 5 Real-Time PCR Instrument (Applied Biosystems, Waltham, MA), with real-time monitoring of amplification progress facilitated by fluorescence data from EvaGreen intercalating dye and ROX reference dye over the 60-minute duration.

#### LAMP assay using oligo-antibody probes with the presence of HIV-1 p24 antigen

Once we determined the starting concentration of the conjugate probes, oligo-conjugated antibody probes were subjected to incubation with HIV-1 p24 antigen at various concentrations prior to their addition to the LAMP master mix. The oligo-antibody probes maintained at a consistent concentration of 10 nM. In the sample volume of five (5) μL, two (2) μL of each oligo-antibody probe 47p112F and 49p125R were mixed in a 1:1 ratio. Subsequently, one (1) μL of HIV-1 p24 antigen at concentrations ranging from 2 fM to 20 nM was introduced to the probe mixture. The probes and antigen mixtures were incubated at room temperature for 30 minutes to ensure immunobinding before mixing with LAMP master mix. SARS-CoV-2 Spike Protein at a concentration of 20 nM served as a non-target antigen to assess the specificity of oligo-conjugated antibody probes towards HIV-1 p24. The LAMP assays were performed at 66°C for 50 minutes. Comparative analysis of the amplification plots for each condition was conducted against an antigen-free condition (47p112F and 49p125R in a 1:1 ratio without antigen). The full-length DNA target of 193 nt at a concentration of 10 nM was used as the positive control DNA, whereas nuclease-free water served as NTC.

### 2.5 ai-LAMP detection limit on LFIA

To allow detection readout on lateral flow strip format, the forward loop primer (LF) and backward loop (LB) primer of LAMP were purchased with fluorescein isothiocyanate (FITC) and biotin labels, respectively. To evaluate the limit of detection of ai-LAMP assay, various concentrations of HIV-1 p24 antigen ranging from 2 fM to 20 nM (1 μL) were incubated with oligo-conjugated antibody probes (2 μL of each probe) at room temperature for 30 minutes to allow immunobinding. The oligo-antibody probes without antibody were used as antigen-free negative control. The two controls for LAMP assays without oligo-antibody probes included nuclease-free water as NTC and full-length 193 nt as positive control. Five (5) μL of samples was added to twenty (20) μL of LAMP mastermix as mentioned above in ai-LAMP assay optimization. The LAMP assays were conducted using conjugate probes with the presence of HIV-1 p24 antigen. The amplicons were then added to commercial Nucleic Acid Probes Detection Devices with a two-test-line format (Ustar Biotechnologies, Hangzhou, China). A negative result is indicated by the absence of any test lines, while a positive result is indicated by the appearance of one or two test lines, depending on the type of labeling molecules used (anti-Digoxigenin or anti-FITC). The test band intensity analysis was performed according to our previously published work [8]. Briefly, after 15 mins of initial sample addition, the LFIAs were scanned using Epson V850 Pro Scanner. A custom MATLAB script created by Holstein et al. [9] was utilized to measure the intensity of the test bands. This script allows for the selection of a region of interest (ROI) around the test line, from which the average grey-scale pixel intensity, referred to as “I_raw_”, is calculated. This raw intensity value is then adjusted by subtracting the average pixel intensity from a local background region, accounting for background interference by using the grey-scale pixel intensity 40 pixels below the test band, termed “I_background_”. The adjusted intensity is then normalized to produce the normalized pixel intensity of the test line, calculated using the formula: I_background_ subtracted = (I_raw_− I_background_) / (0 − I_background_), where 0 represents the minimum possible pixel intensity. The limit of detection (LOD) was assessed using a one-way ANOVA with Dunnett’s post hoc test for multiple comparisons, performed with GraphPad Prism (GraphPad Software, Boston, MA), comparing the LFIA test bands at each concentration against the negative controls (without template) with a 95% confidence interval.

## 3. Results and discussion

### 3.1 Overview of ai-LAMP concept

We developed a novel HIV-1 p24 protein detection assay that provides a straightforward, visual readout in the same manner as a conventional RDTs. While LFIA tests for p24 have been developed, Stone et al. conducted a multi-laboratory comparison of laboratory-based p24 Ag assays and Ag RDTs [10]. They found that the RDTs were unable to detect HIV from a single clinical sample even at viral loads up to 1,000,000 vp/mL (approx. 3.76 pM p24) [10]. Another study by Livant et al. and their colleagues revealed that p24 Ag/Ab tests have been updated and provide improved detection, identifying between 28% and 64% of acute HIV infections [11]. Additionally, while it has been noted that minor sequence variations can impact the tertiary structure of p24, these alterations do not affect the sensitivity of assays based on p24 antigen detection. This is an advantage of p24 antigen detection, as it remains effective across a variety of HIV subtypes compared to RNA-based detection [12]. Like conventional RDTs, our assay targets the HIV-1 p24 antigen, a key capsid protein indicative of HIV-1 infection and viral load, given its presence in approximately 2,000-3,000 copies per virion [13]. Unlike conventional RDTs, which target p24 directly, we employ LAMP to generate billions of copies of target oligonucleotides upon the binding between p24 and its antibody. This amplifies the signal and enables detection through LFIA.

The assay involves the development of ai-LAMP, as illustrated in Schematic 1. Two monoclonal antibodies specific to HIV-1 p24 are conjugated with partial DNA targets for the LAMP reaction. These partial targets feature a short overlapping region of six nucleotides, facilitating their hybridization to form a complete template for LAMP. Alternative pairings of partial targets with longer overlapping regions of 9, 12, and 15 nucleotides were also evaluated; however, the six-nucleotide overlap was selected as optimal, as it minimized nonspecific background hybridization and thereby preventing unintended LAMP initiation in the absence of antigen-probe binding. Upon the presence of HIV-1 p24 antigens in a sample, binding to the antibody probes brings the partial DNA targets into close proximity, accelerating theirhybridization into a full-length template for the LAMP reaction. Consequently, LAMP amplification occurs much more rapidly than without antigen present. We exploit the differential amplification times of the LAMP reaction with and without HIV-1 p24 antigens to provide a visual readout on lateral flow strips. For LFIA readout, the amplicons are doubly labeled with fluorescein isothiocyanate (FITC) and biotin during amplification via the incorporation of labeled loop primers. FITC facilitates attachment of amplicons to the LFA strip surface via an anti-FITC antibody on the test line, while biotin serves as a capture moiety for streptavidin immobilized on gold nanoparticles (AuNPs).

**Schematic 1.**
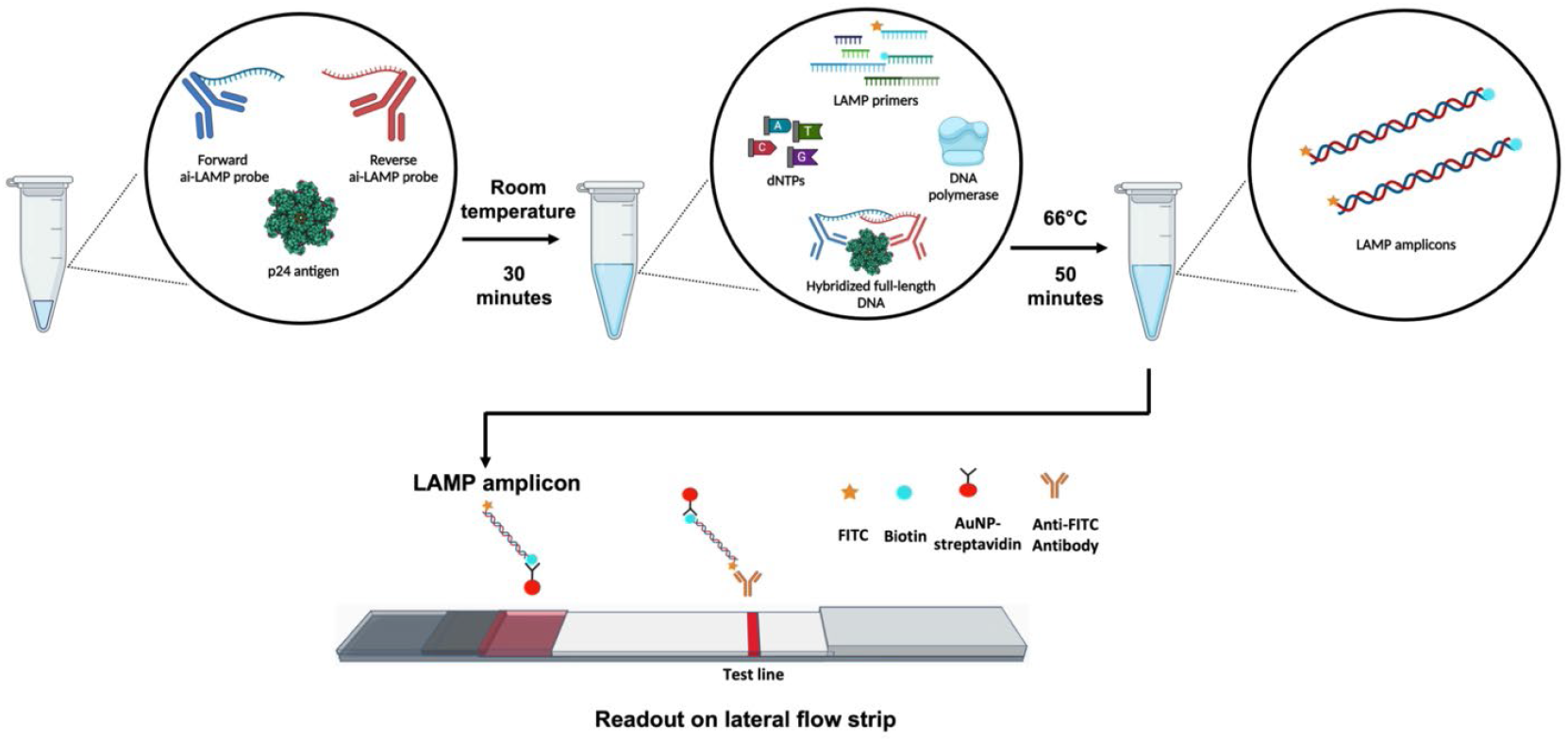
ai-LAMP assay workflow depicting the detection principle of the ai-LAMP assay. First, two forward and reverse ai-LAMP probes (p24 monoclonal antibodies conjugated to partial DNA targets) were incubated with HIV-1 p24 antigens at room temperature for 30 minutes. Subsequently, a LAMP mastermix was added to the tube. The reaction mixture was then incubated at 66°C for 50 minutes to generate amplicons. For visual readout, the LAMP amplicons, labeled with FITC and biotin molecules, were introduced onto the test strip. Gold nanoparticles coated with streptavidin (AuNP-streptavidin) then bind to the biotin in the LAMP product, forming a complex that migrates through the test strip due to capillary action. The appearance of a test line on the strip indicates the successful detection of the LAMP product by AuNP-streptavidin, which has been captured by the FITC-antibody (Anti-FITC Ab) on the test line. The cartoon p24 structure was constructed based on PDB 4XFX [14].

### 3.2 ai-LAMP primer and template design

In this work, the LAMP reaction serves as a signal amplification upon the binding between HIV p24 antigen and its paired antibodies. Therefore, to avoid false positive amplification without the presence of HIV p24 antigen, the primer candidates should be orthogonal to HIV and human genomes and should not be found in blood samples. We selected the primer set targeting cholera enterotoxin subunit A *(ctxA)* gene of *Vibrio cholerae* (*V. cholerae*) (Table S1). The NCBI BLAST confirmed that there is no match between selected primers against human and HIV genomes. The partial forward (F) and reverse DNA target (R) with 6 nucleotide overlapping region were displayed in Table S2. *V. cholerae* is a waterborne pathogen that causes gastrointestinal illness but does not circulate in the blood. One case report has noted instances of *V. cholerae* inducing bacteremia, however, the incidence of *V. cholerae* in blood is negligible [15].

The amplification plots in Figure 1A and B show that the LAMP assays with the presence of either F or R with various concentrations did not amplify during the 1-hour incubation time. On the other hand, as seen in Figure 1C, the samples with combinations of F and R at 100 nM, 10 nM and 1 nM began to amplify approximately at 25, 35, and 45 minutes, respectively. The full-length positive control at 10 nM started to amplify early within 10 minutes. We hypothesized that the amplification of both DNA probes (FR) was slower than that of the full-length DNA target because the probes need to locate and hybridize with each other to form a complete template. The NTC samples showed no amplification, confirming that this primer set did not cause primer-dimers which could lead to false-positives. Taken together, the selected primers and designed targets could serve as signal amplification for the ai-LAMP assay.

**Figure 1.**
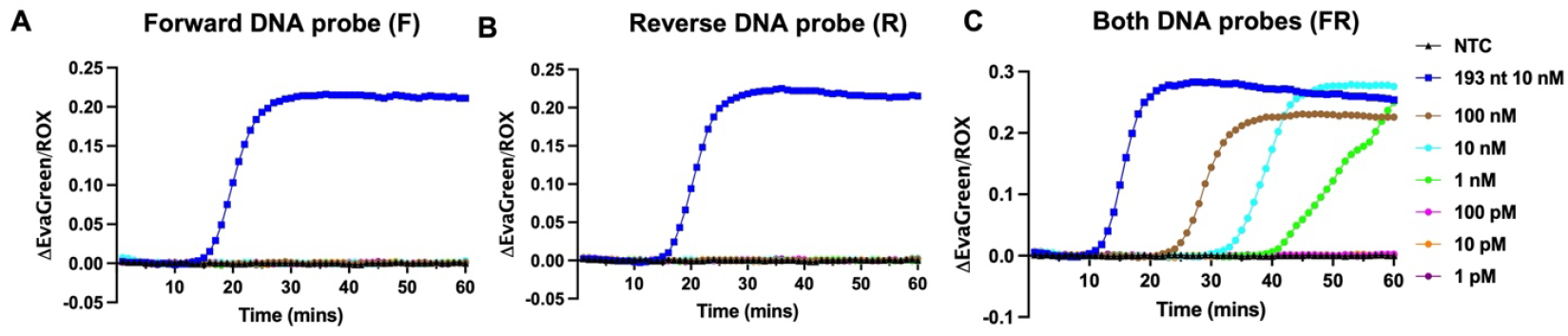
ai-LAMP primer and template design. Representative amplification plots of LAMP reactions with (A) the forward DNA target of 112 nt (F), (B) the reverse DNA target of 125 nt (R), and (C) combinations of the forward and reverse targets (FR). N = 2. Another replicate is shown in Figure S1. No antigen is added in these reactions.

This experiment suggests that for the LAMP reaction to proceed effectively, the presence of both F and R at specific concentrations is essential. Both the forward and reverse LAMP sequences with small overlapping region are essential to produce the full-length target for successful LAMP reactions. Further, the concentrations of F and R DNA in DNA-antibody probes are critical in the LAMP initiation.

### 3.3 Analysis of oligo-conjugated antibody probes

The selection of antibody pairs specific to HIV-1 p24 is crucial for the design of our assay. It is essential that the interaction between the antibodies and the p24 antigen be both specific and rapid to form a robust antigen-antibody complex. A pair of monoclonal antibodies, mAB247p and mAB249p, demonstrated high binding affinity and low equilibrium dissociation constant (KD) values, as assessed by Octet RED96 Biolayer Interferometry (Forte Bio, Menlo Park, CA) [16]. Consequently, mAB247p and mAB249p were selected as the antibody probes for the development of the ai-LAMP assay.

Copper-free click chemistry was utilized as a method for conjugating antibodies and oligonucleotides. This method is based on the strain-promoted alkyne-azide cycloaddition (SPAAC) reaction between a DBCO moiety and an azide-labeled partner. The protein-stained gels of 47pF and 49pR in Figure 2A and C exhibited distinct band patterns for the azide-activated antibodies in lane 2 and for the oligo-antibody conjugates in lane 3. The azide-activated antibody displayed a 150 kDa band, corresponding to the molecular weight of an intact mouse IgG. The oligo-conjugated antibodies in lane 3 revealed shifted bands above 150 kDa due to the formation of higher molecular weight complexes with conjugated oligonucleotides. The estimated sizes of the oligo-conjugated antibodies, with one oligo per antibody, were 185 kDa for 47pF and 188 kDa for 49pR, respectively, which is consistent with the observed products on the gels. Additionally, faint bands above 250 kDa were observed in both conjugates, corresponding to the presence of three oligos per antibody. For three oligos per antibody, the estimated sizes were 256 kDa for 47pF and 267 kDa for 49pR. It’s worth noting that we observed fragment bands ranging from 75 to 100 kDa and a distinct band at 50 kDa (Figure S2) after conjugation. These bands corresponded to non-intact antibody in various forms such as two heavy chains, one light chain and one heavy chain, and one heavy chain, which appear at 100 kDa, 75 kDa, and 50 kDa, respectively [17].

**Figure 2.**
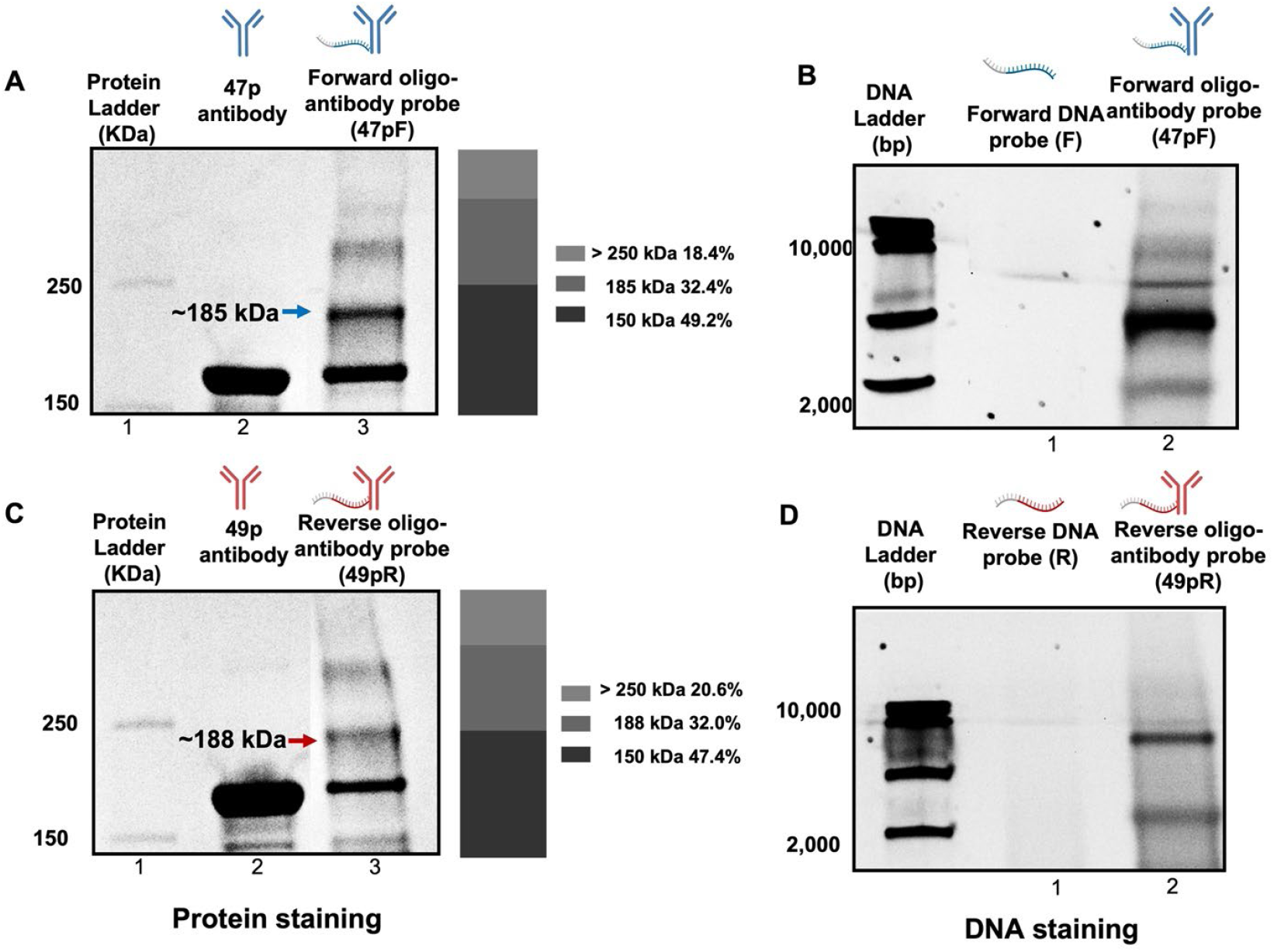
The SDS-PAGE gel analysis of oligo-antibody probes. (A) The protein staining of azide-modified 47p antibody and forward 47pF conjugates. (B) DNA staining of DBCO-modified forward DNA probe F and forward 47pF conjugates. (C) The protein staining of azide-modified 49p antibody and reverse 49pR conjugates. (D) DNA staining of DBCO-modified reverse DNA probe R and reverse 49pR conjugates. The bar charts next to protein staining gels represent the percentage yield of each band in lane 3.

DNA-stained gels of oligo-antibody conjugates are shown in Figure 2 B and D for lane 3. The DBCO-modified F and R oligos have approximate molecular weights of 35 kDa and 39 kDa, respectively, and appeared as major bands between 150 and 300 bp (Figure S3), which were omitted in lane 2 of Figure 2B and D. Overlaying the gel images of protein-stained and DNA-stained gels revealed that these bands corresponded to the positions of protein bands between 150 kDa and 250 kDa, as well as the smear bands above 250 kDa, with intense bands appearing at 185 kDa and 188 kDa. We found that SYBR Gold staining could potentially stain protein products; this phenomenon has also been observed by others [18].

To avoid the confounding effect of protein staining by SYBR Gold, protein-stained gel images were used to analyze conjugate band intensity and calculate conjugation yield through ImageJ software. The bar charts next to the protein-stained gels (Figure 2A and C) represent the percentage yield of each band in lane 3. For 47pF conjugates, the yield of the 185 kDa product was 32.4%, and the yield of the band above 250 kDa was 18.4%. For 49pR conjugates, the yield of the 188 kDa product was 32.0%, and the yield of the band above 250 kDa was 20.6%.

A limitation of the work is the relatively inefficient conjugation of the oligo-antibody reaction. In initial conjugation studies, we explored strategies that could improve conjugation efficiency. The following parameters, including the number of azide molecules on the antibody, the molar ratio of functionalized antibody to oligonucleotide, the conjugation time, and the reaction temperature, are crucial for achieving high yields of the desired click reaction products [6]. Further work could include pre-treating DNA probes with betaine or DMSO to reduce the formation of secondary structures before subjecting to bioconjugation to improve conjugation efficiency. The conjugated DNAs used in this work were relatively long as compared to other publications in the antibody oligo conjugation works and are likely to possess secondary structures and hairpin structures, which could have led to hindrance of functional group for click chemistry reaction [19]. Ultimately, the limited conjugation efficiency could be mitigated through outsourcing the conjugation now that numerous companies have emerged to create oligo-antibody conjugates.

### 3.4 ai-LAMP proof-of-concept

#### LAMP assay using oligo-conjugated antibody probes without HIV-1 p24 antigen

To assess potential non-specific amplification induced by the partial probe, LAMP assays were performed using either of the oligo-conjugated antibody 47pF or 49pR probes at various concentrations. Analysis of the amplification plots shown in Figure 3A and B revealed that neither 47pF nor 49pR alone caused non-specific amplification at any tested concentration. Only reactions with the positive control DNA (full-length DNA target) produced amplification. This finding is crucial because the LAMP reaction is used for signal amplification in this ai-LAMP assay. It is essential that the partial probes do not trigger the LAMP reaction, as this could result in false-positive outcomes.

**Figure 3.**
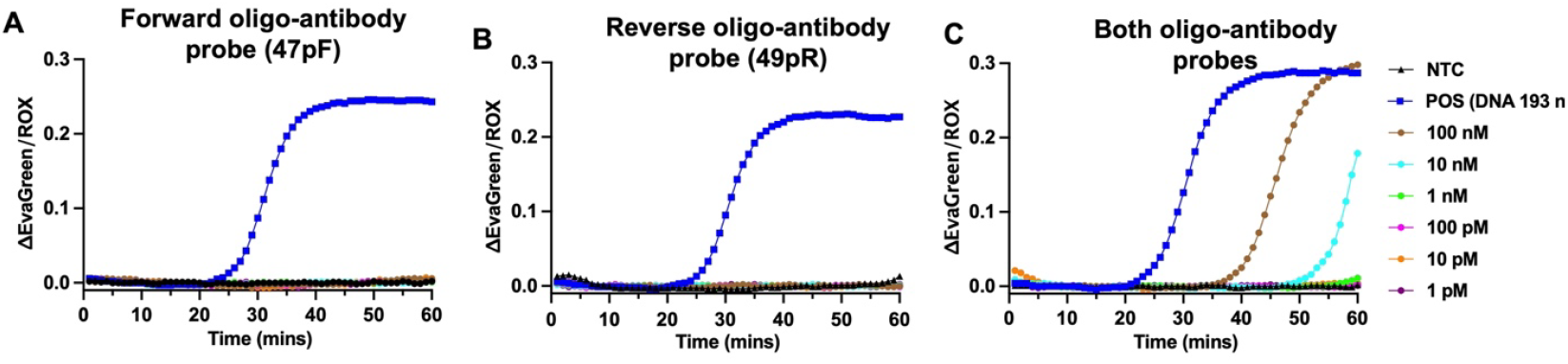
LAMP assays using oligo-antibody probes without HIV-1 p24 antigen. The ctxA LAMP assay using 47pF probes only (A), 49pR probes only, and (C) the combinations of both probes at various probe concentrations as shown.

Following the evaluation of partial oligo-antibody probes, LAMP assays were conducted using a mixture of both probes, 47pF and 49pR, at concentrations ranging from 1 pM to 100 nM. The probes were combined in a 1:1 ratio, and full-length DNA template and nuclease-free water served as positive and negative controls, respectively. The amplification plot results illustrated in Figure 3C and Figure S4 demonstrated that the probe mixture at concentration of 100 nM and 10 nM exhibited amplification within 60 minutes. At lower probe concentrations, it was anticipated that the LAMP assay would show amplification with both oligo-antibody probes even in the absence of antigen, as in Figure 1. Given adequate incubation time and optimal temperature, both conjugated oligonucleotide targets are expected to hybridize and create a full-length target suitable for the LAMP assay. However, this time exceeded our 60-minute cutoff, and no detectable amplification occurred at concentrations below 1 nM, likely due to steric hindrance caused by the bound antibodies reducing DNA hybridization. For the next steps, probes at a concentration of 10 nM were chosen, as this concentration consistently demonstrated the onset of amplification towards the end of the amplification period.

#### LAMP assay using oligo-conjugated antibody probes with the presence of HIV-1 p24 antigen

The oligo-conjugated antibody probes at a concentration of 10 nM were selected based on the observation that at this concentration, we only detect amplification without antigen late in the assay around 55 minutes. Our hypothesis was that the presence of HIV-1 p24 antigen would expedite the binding and hybridization of the two partial oligonucleotides, 112F and 125R, leading to accelerated amplification compared to conditions without antigen, creating a marked (left) shift in the amplification curves. As shown in the amplification plot in Figure 4A, in which the amplification period was set at 50 minutes, the LAMP reactions with HIV-1 p24 antigen were amplified within 35 to 45 minutes, whereas the antigen-free conditions did not show amplification. Compared to the amplification profile of oligo-antibody probes without antigen shown in Figure 3C, LAMP reactions with oligo-antibody probes in the presence of HIV-1 p24 antigen exhibited faster amplification by approximately 10 to 20 minutes in a concentration-dependent manner Analysis of Ct values revealed statistically significant differences in Ct values of LAMP reactions when using oligo-conjugated antibody probes with and without antigens as samples (Figure S5). To verify that the accelerated amplification resulted from the binding of HIV-1 p24 antigen to the antibody probes, which facilitates the proximity of the oligo-antibody probes, SARS-CoV-2 Spike protein was used as a non-target antigen and incubated with the oligo-antibody probes. As shown in Figure 4A, the LAMP reaction was not triggered by the presence of the Spike protein, even at the highest concentration of 20 nM. This result indicates that the shift in amplification time was due to the specific binding of HIV-1 p24 antigen to its monoclonal antibodies, 47p and 49p.

**Figure 4.**
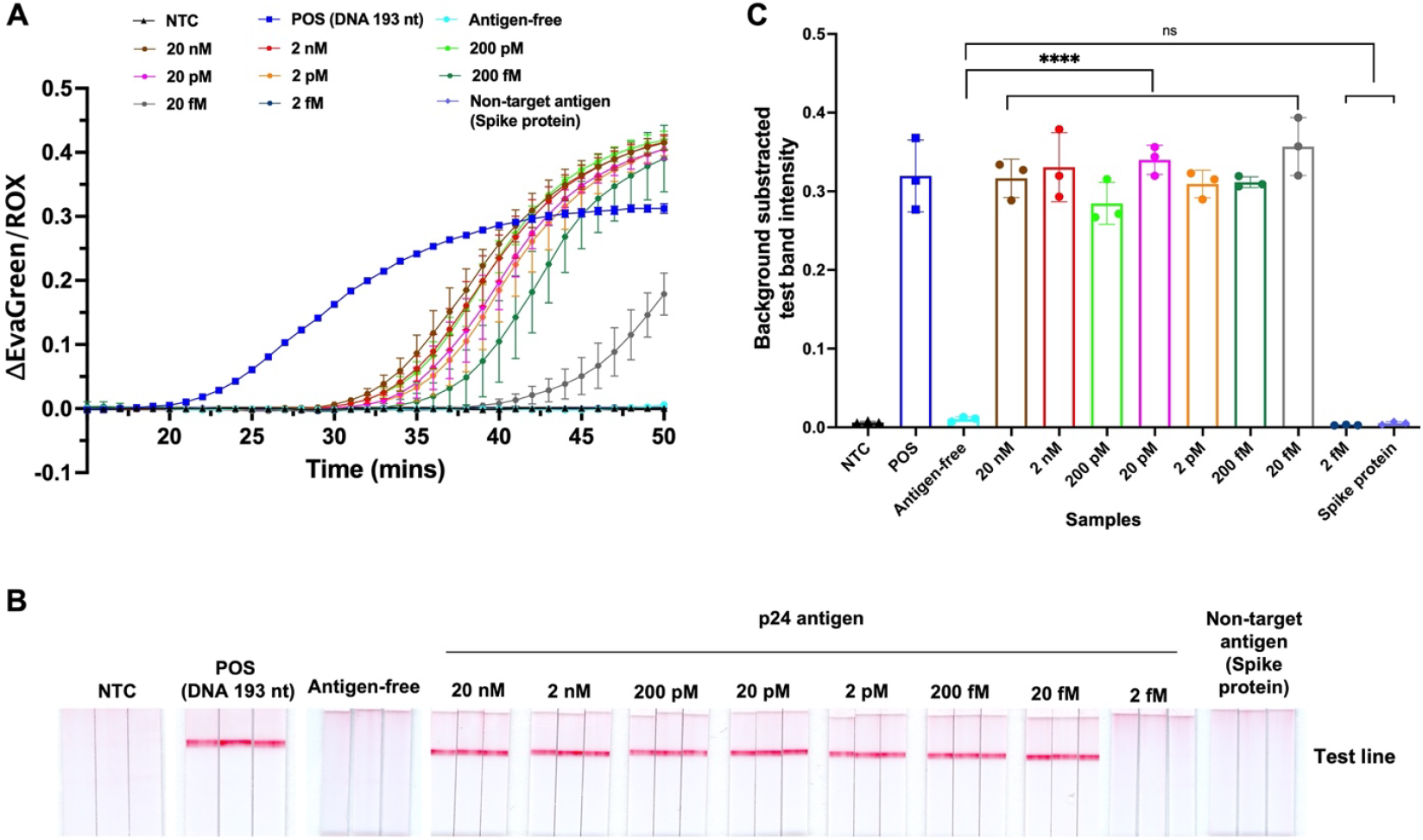
LAMP assay using oligo-conjugated antibody probes in the presence of HIV-1 p24 antigen. (A) The amplification plot of LAMP assays using both oligo-antibody probes with various concentrations of HIV-1 p24 antigen compared to non-amplifying conditions: NTC, antigen-free, and SARS-CoV-2 Spike protein as a non-target antigen. (B) ai-LAMP assay readout on the LFIA platform. The amplicons from LAMP assays using both oligo-antibody probes incubated with various concentrations of HIV-1 p24 and non-target antigen are visualized on LFIA strips. C) Corresponding test band intensity analysis of LFIA strips shown in B. n = 3; **** indicates p ≤ 0.0001.

Taken together, these experiments demonstrate the proof-of-concept for the ability of the ai-LAMP assay to detect HIV-1 p24 through the specific binding of HIV-1 p24 to its corresponding antibodies, combined with the signal amplification provided by LAMP. The designed partial DNA targets successfully conjugated with monoclonal antibodies specific to HIV-1 p24, serving as templates for the LAMP reaction. In the presence of HIV-1 p24 antigen, the amplification time of LAMP reactions was expedited due to the binding of HIV-1 p24 antigens to antibody probes. This interaction facilitated the proximity of two partial DNA probes conjugated to antibodies, thereby expediting the hybridization of partial DNA probes to form the full LAMP target.

Of note, all experiments in this section were performed using the oligo-conjugated antibodies from the same batch. While the conjugation efficiency was 32 % for 186 -188 kDa products and 18 – 20 % for product above 250 kDa, we can see the discrepancy in time to amplification and Ct values of LAMP reactions between samples with and without HIV-1 p24 antigens. We expect that the improvement of conjugation efficiency of oligo-conjugated probes would allow us to achieve improved detection sensitivity and attain dynamic ranges of the ai-LAMP assay. Assay optimizations that could be further investigated include immunobinding buffer and time as these two factors affect the binding kinetic between antigen and antibody [20]. Such optimizations may be especially important when adapting ai-LAMP for detecting low-abundance p24 antigen in complex matrices like plasma or whole blood, where immune complexes and matrix effects may interfere. In addition, Zhang et al. reported that increasing sample volume could improve the detection sensitivity of proximity extension assay, which is a similar detection platform that used an oligo-conjugated antibody as a probe [2].

### 3.5 The limit of detection (LOD) on lateral flow strips

LFIAs are commonly employed for colorimetric readouts in various viral Rapid Diagnostic Tests (RDTs) due to their user-friendly nature. Typically, results obtained through LFIA are qualitative, providing a binary outcome where one line indicates a negative result, and two lines indicate a positive result. However, quantitative data can be attained by converting test line intensities into numerical values using computer software [21]. To enable LFIA readouts, loop primers LF and LB were labeled with FITC and biotin, respectively, and incorporated into amplicons following amplification. LAMP assays using two oligo-conjugated probes, 47pF and 49pR, incubated with HIV-1 p24 concentrations ranging from 2 fM to 20 nM, were conducted. The amplicons were visualized on LFIAs to determine the Limit of Detection (LOD) by quantifying test band intensity. Unlike commonly used LFIA with one control line and one test line format, the amplicons were added to LFIA device with two test line-format, where no test lines represent negative result while one or two test lines represent positive result depending on type of labeling molecules (anti-Digoxigenin, anti-FITC). This LFIA device was chosen over the standard one control line and one test line format due to its superior performance in detecting amplicons and displaying a test line for low concentration antigens, which the regular LFIA format may only show as a faint test band [22].

The FITC test lines were visible in LAMP assays with HIV-1 p24 concentrations ranging from 20 fM to 20 nM and positive control DNA due to the FITC labeled LF incorporation. In conversely, the NTC sample, antigen-free condition, and Spike protein samples did not show any lines. The LOD of the assay was established by quantifying test band intensity using a custom MATLAB script, which calculated the average grayscale pixel intensity of the test band and subtracted the average background pixel intensity as shown in Figure 4C. Statistical analysis using one-way ANOVA with Dunnett’s post hoc revealed a significant difference in test band intensity for HIV-1 p24 concentrations ranging from 20 fM to 20 nM compared to the antigen-free condition. According to this LFIA readout, the ai-LAMP assay could detect HIV-1 p24 down to 20 fM, which is equivalent to 532 fg/mL of p24. It’s worth noting that the readout time was set to 15 minutes for image processing consistency. However, the test line can be visualized within 5 minutes upon adding samples to the test strips. Additionally, acceptability of this increased assay time compared to rapid Ag tests will be an important consideration for implementation in POC settings. The total test time from antigen-probe incubation to result readout by naked eyes is 85 minutes which is less time than most of laboratory-based nucleic acid and Ab/Ag tests [23].Two steps that could be further optimized to reduce total assay time are the sample and probes incubation times given that p24 Ag/Ab binding can occur within seconds for existing RDTs. Amplification speed can be improved by using an alternative DNA polymerase with enhanced performance, for example, OmniAmp pol can effectively amplify DNA or RNA templates in just 30 minutes, which is 20% quicker than conventional LAMP methods [24].

In individuals with acute HIV infection, the concentration of p24 antigen in the blood can be as high as several thousand pg/mL. However, reliable detection of low concentrations of p24 antigen, especially during the very early stages of HIV infection is critical for early treatment and transmission prevention. The primary challenge associated with acute HIV detection using RDTs is the wide clinical range of p24 antigen concentrations found in the blood. This range extends across at least four orders of magnitude, from under 1 to over 10^3^ pg/mL. [25] The ai-LAMP assay presented in this work has a dynamic range of 0.5 to 5 x 10^5^ pg/mL. As of July 2025, the Determine™ HIV–1/2 Ag/Ab Combo is the only FDA-approved POC assay in lateral flow assay format, with a claimed LOD of 25 pg/mL [26]. However, the assay has faced challenges related to variable performance, sensitivity issues during acute infection, and specificity concerns in clinical testing [25].

A limitation of our test is that the experiments were validated using purified HIV-1 p24 antigen. Components in biological samples such as plasma and whole blood could reduce antibody-binding or interfere with the LAMP assay, leading to reduced sensitivity. For p24, in particular, immune complex disruption via heat or acid treatment may be required for ai-LAMP to compete with host antibodies during the chronic infection stage [25]. However, many existing sample preparation methods exist to mitigate these challenges.

## 4. Conclusion

This study presents a proof-of-concept for a novel signal amplification, ai-LAMP, for protein detection with an LFIA detection readout. This protein detection assay can be adapted to other proteins by changing the antibody-moiety. Here, ai-LAMP uses monoclonal antibody probes specific to HIV-1 p24 conjugated with partial overlapping DNA LAMP targets to demonstrate detection of p24. Selecting the optimal concentration of oligo-conjugated probes is crucial as it directly impacts the rate of amplification. For HIV-1 p24 ai-LAMP, an oligo-conjugated probe concentration of 10 nM, mixed in a 1:1 ratio, provided consistent amplification performance. At this concentration, we observed a statistically significant difference in the time to amplification of the LAMP assay with, and without, the presence of p24 antigen. Antigen binding brought the partial DNA templates conjugated each of the antibodies into proximity with one another, expediting hybridization and subsequent LAMP amplification, which we utilized for visual detection via LFIA. Specificity analysis using SARS-CoV-2 spike protein as a non-target antigen revealed that the shift in amplification time occurred only with HIV-1 p24 as the target. The assay workflow comprised 30 minutes of immunobinding between the oligo-antibody probes and p24 antigen, followed by 50 minutes of amplification at 66°C, and a 5–15-minute readout on LFIA for a total assay time of 85-90 minutes. The ai-LAMP assay currently achieves an LOD of 532 fg/mL, which represents a more than 46-fold improvement over commercial p24 antigen RDTs. Overall, the novel ai-LAMP assay shows potential for enhancing protein-based detection with simple LFIA readout. This platform does not require sophisticated laboratory equipment, thus enabling decentralization of the test to point-of-use locations.

