## Supplemental Information for "Antibody-Initiated Loop-Mediated Isothermal Nucleic Acid Amplification (ai-LAMP) as a New Biosensor for Antigen Detection" (from another version of this preprint: version 1, 2025-09-24)

### Supporting Information

| **Nucleotide sequences of LAMP primers that target the *ctxA*** | |
|---|---|
| Primer | Sequence (5’ – 3’) |
| F3 | TCGGGCAGATTCTAGACC |
| B3 | GTGGGCACTTCTCAAACT |
| FIP | TTGAGTACCTCGGTCAAAGTACT-TCCTGATGAAATAAAGCAGTCA |
| BIP | TCAACCTTTATGATCATGCAAGAGG-GGAAACATATCCATCATCGTG |
| LF-FITC | /56-FAM/ GGTGGTCTTATGCCAAGAGG |
| LB-Bio | /5BiosG/CCTAACAAATCCCGTCTGAGTT |

Table S 1 LAMP primers targeting ctxA gene of Vibrio Cholerae

| Target | Size (nt) | Sequence (5’ – 3’) |
|---|---|---|
| Full-length | 193 | TCGGGCAGATTCTAGACCTCCTGATGAAATAAAGCAGTCAGGTGGTCTTATGCCAAGAGGACAGAGTGAGTACTTTGACCGAGGTACTCAAATGAATATCAACCTTTATGATCATGCAAGAGGAACTCAGACGGGATTTGTTAGGCACGATGATGGATATGTTTCCACCTCAATTAGTTTGAGAAGTGCCCAC |
| Forward (F) | 112 | /Amine/TGATGATAAGTTATATCGGGCAGATTCTAGACCTCCTGATGAAATAAAGCAGTCAGGTGGTCTTATGCCAAGAGGACAGAGTGAGTACTTTGACCGAGGTACTCAAGGATAG |
| Reverse (R) | 125 | /Amine/AGTTTGACCCACTAAGTGGGCACTTCTCAAACTAATTGAGGTGGAAACATATCCATCATCGTGCCTAACAAATCCCGTCTGAGTTCCTCTTGCATGATCATAAAGGTTGAGTAGAGCGTCTATCC |

Table S 2 Designed LAMP targets (GenBank: NC_002505.1) including the full-length target of 193 nt as a positive control, forward and reverse templates with 6 nucleotides overlapping (underline) allowing the hybridization to form full-length target upon the proximity contact of corresponding antibodies. Note: Amine is 5AmMC6 modification from IDT.

| **Reagent** | **Final Concentration** | **Actual Rxn (25µLVol)** | **Master Mix (1.1 x # Rxns x Aliquot Vol)** |
|---|---|---|---|
| Nuclease free Water | ---- | 13.5 | 118.80 |
| Isothermal buffer I | 1x | 2.5 | 22.00 |
| dNTPs (100 mM) | 1 mM | 0.25 | 2.20 |
| EvaGreen (20X) | 0.2X | 0.25 | 2.20 |
| ROX Dye (50X) | 1X | 0.5 | 4.40 |
| Primer mix 10x | 1x | 2.5 | 22.00 |
| Bst DNA polymerase 2.0 (8U/µL) | 4U | 0.5 | 4.40 |
| **Mastermix** | | **20** | |
| **Template** | ---- | 5 |  |
| **Total** | | **25** | |

Table S 3 LAMP assay components for ai-LAMP

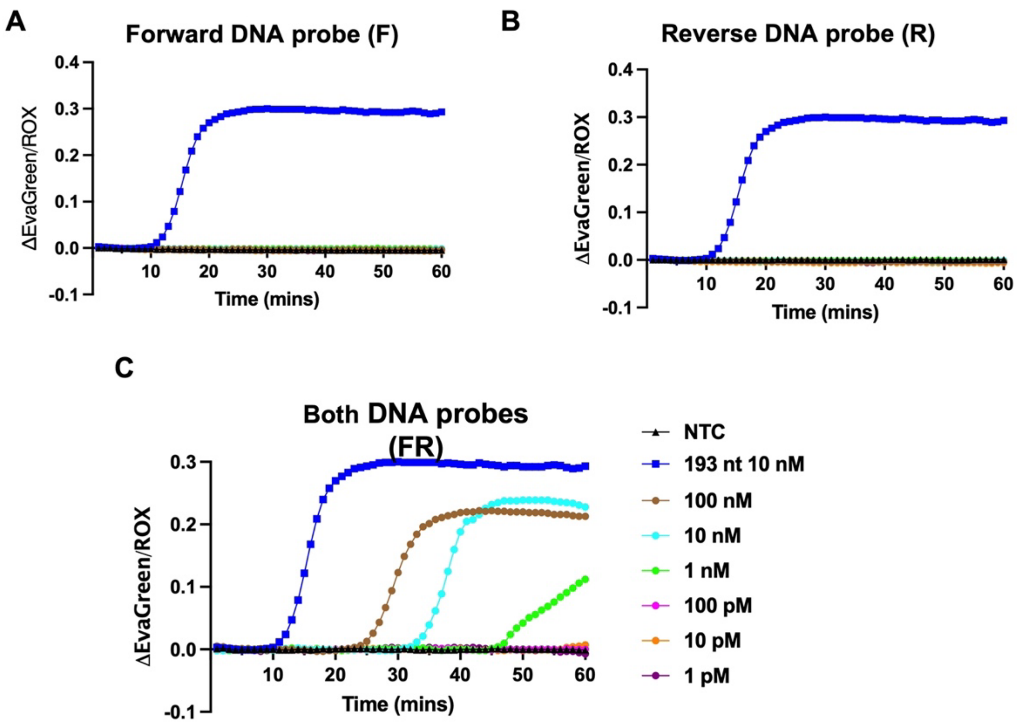

Figure S 1. Replicate 2 of LAMP reactions with (A) forward DNA target of 112 nt (F), (B) reverse DNA target of 125 nt (R), and (C) combinations of forward and reverse targets (FR).

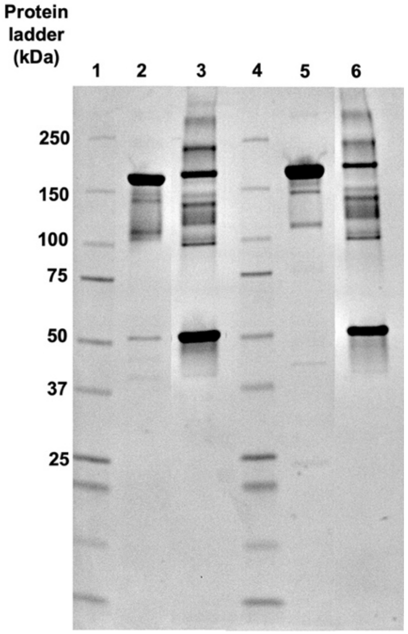


Figure S 2. Non-reducing SDS-PAGE gel analysis of oligo-conjugated antibody probes. The protein staining of azide-modified 47p antibody (lane 2), forward 47pF conjugates (lane3), azide-modified 49p antibody (lane 5), and reverse 49pR conjugates (lane 6)

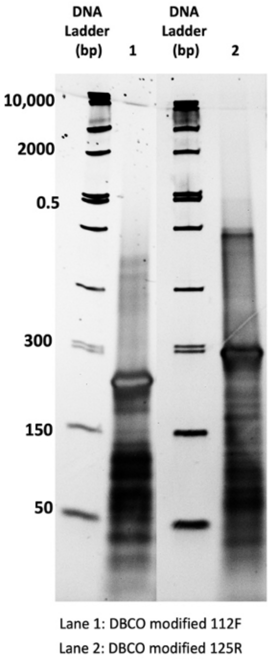


| | Nucleotide length (nt) | MW with modification (g/mol) | Concentration (μmol/L) |
|---|---|---|---|
| 112F_DBCO | 112 | 35657.48 | 15.3 |
| 125R_DBCO | 125 | 39281.78 | 15.3 |

Figure S 3. SYBR gold staining-gel of DBCO-modified F and R and concentration of DBCO modified DNA probes.

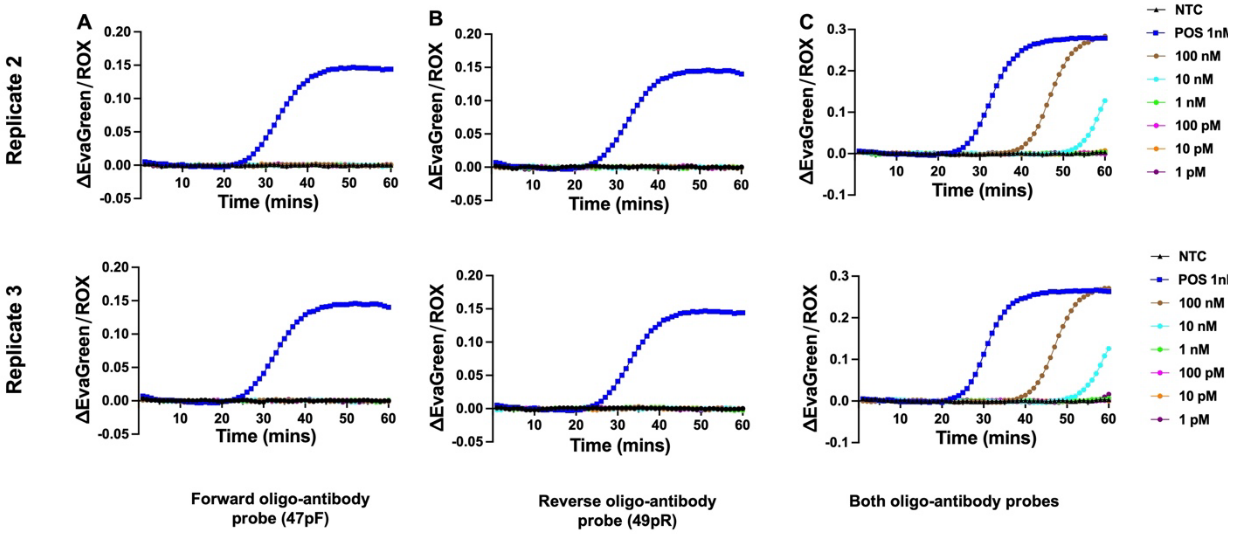


Figure S 4. Replicate 2 and 3 of LAMP reactions with (A) forward oligo-antibody probe, (B) reverse oligo-antibody probe, and (C) combinations of both probes.

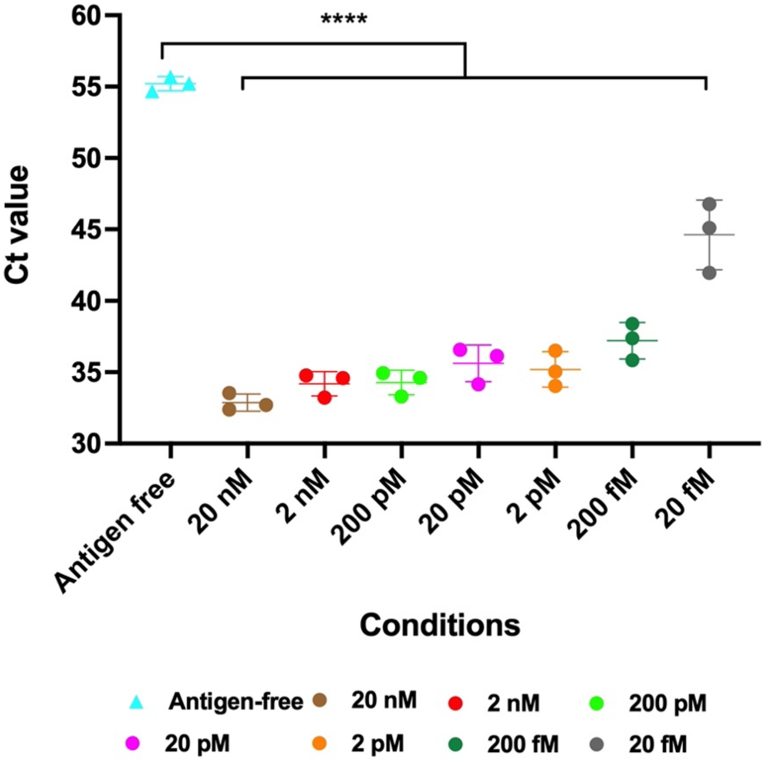


Figure S 5. The Ct value of ai-LAMP assays using both oligo-antibody probes with the presence of various concentrations of HIV-1 p24 antigen as compared to antigen-free condition.

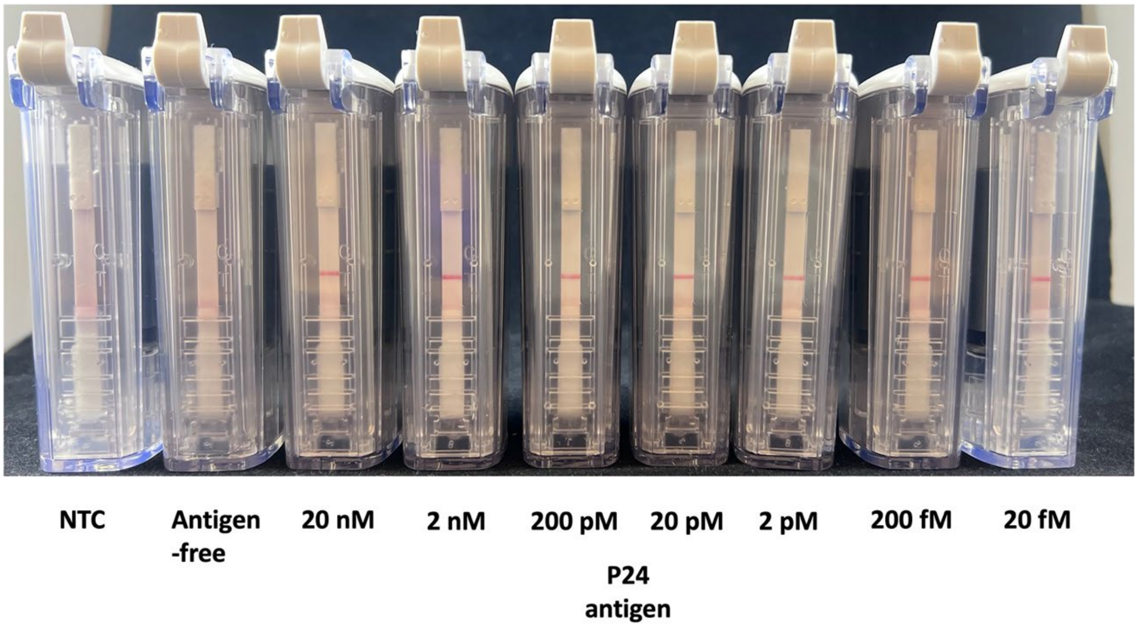


Figure S 6. LFIA device used in this work as the detection unit. The LFIA strips with one line indicate a positive result while no line represents a negative result.
